# Ion Mobility for Unknown Metabolite Identification: Hope or Hype?

**DOI:** 10.1101/2022.08.26.505158

**Authors:** Carter K. Asef, Markace Rainey, Brianna M. Garcia, Goncalo J. Gouveia, Amanda O. Shaver, Franklin E. Leach, Allison M. Morse, Arthur S. Edison, Lauren M. McIntyre, Facundo M. Fernández

## Abstract

Ion mobility (IM) spectrometry provides semi-orthogonal data to mass spectrometry (MS), showing promise for identifying unknown metabolites in complex non-targeted metabolomics datasets. While current literature has showcased IM-MS for identifying unknowns under near ideal circumstances, less work has been conducted to evaluate the performance of this approach in metabolomics studies involving highly complex samples with difficult matrices. Here, we present a workflow incorporating *de novo* molecular formula annotation and MS/MS structure elucidation using SIRIUS 4 with experimental IM collision cross-section (CCS) measurements and machine learning CCS predictions to identify differential unknown metabolites in mutant strains of *Caenorhabditis elegans*. For many of those ion features this workflow enabled the successful filtering of candidate structures generated by *in silico* MS/MS predictions, though in some cases annotations were challenged by significant hurdles in instrumentation performance and data analysis. While for 37% of differential features we were able to successfully collect both MS/MS and CCS data, fewer than half of these features benefited from a reduction in the number of possible candidate structures using CCS filtering due to poor matching of the machine learning training sets, limited accuracy of experimental and predicted CCS values, and lack of candidate structures resulting from the MS/MS data. When using a CCS error cutoff of ±3%, an average 28% of candidate structures could be successfully filtered. Herein, we identify and describe the bottlenecks and limitations associated with the identification of unknowns in non-targeted metabolomics using IM-MS to focus and provide insight on areas requiring further improvement.

---

Liquid-chromatography mass-spectrometry (LC-MS) remains a powerful technique for interrogating perturbations of small molecules in living organisms through non-targeted metabolomics studies. To relate detected ion features to biological processes, annotation of such features must be performed to assign structure. This is typically accomplished by collecting MS/MS spectra for features of interest and searching against large databases of previously characterized or computationally predicted MS/MS databases such as the Human Metabolome Database^1^, LIPID MAPS structure database^2^, and MassBank^3^. For well-characterized cell lines and organisms, database searches will typically annotate 10% or less of measured features as the coverage of such databases is still incomplete^4^. Annotation of this remaining “metabolic dark matter” is a lengthy process and remains a critical bottleneck in the field of metabolomics^5^.

Limited structural information can be gleaned from the fragment masses of a small molecule’s product ion MS/MS spectrum, though full structural elucidation is not often possible with this information alone. Computed fragmentation trees (*e.g*. from graph theory) have been proposed as a solution to leverage this limited information, providing likely substructures in the unknown molecules^6^. These trees are visualized as a graph of masses and their related molecular formulas showing the logical progression from precursor to product ion, neutral and radical losses, and subsequent products resulting from multiple fragmentation pathways. CSI:FingerID is an *in silico* machine learning (ML) method that successfully extracts structural motifs (or “molecular fingerprints”) from fragmentation trees, which can then be compared against large databases of molecules without archived MS/MS data^7^. This allows for the molecular fingerprint of an unknown feature to be compared against the millions of entries in molecular structure databases such as PubChem to produce a list of ranked candidate structures. This approach has shown promise in the critical assessment of small molecule identification (CASMI) contests as a part of the SIRIUS 4^8^ suite of compound identification tools where it was able to correctly identify 74.8% of features in the top 10 ranked candidates for positive ion mode LC-MS/MS data^8^. Several other computational methods have been proposed that similarly offer a large advantage over manually reviewing MS/MS data, though all generate large lists of candidates that may or may not contain the correct structure in the top ranks^9^.

Ion mobility (IM) spectrometry readily pairs with LC-MS analysis (LC-IM-MS) and offers semi-orthogonal information in the form of collision cross section (CCS) values, a two-dimensional, rotationally averaged representation of molecular size and shape. These CCS values can be derived experimentally from the drift time of an ion feature in the case of drift tube IM (DTIM) or can be calibrated from known DTIM CCS values in the case of traveling wave IM (TWIM)^10^. CCS databases have been developed recently along with machine learning (ML) tools that leverage these experimental values to predict the CCS of molecular structures of interest^11–13^. As CCS is an intrinsic trait of a given molecule, it has been touted as a means to further filter candidate structures generated by *in silico* MS/MS analysis software. ML can then be used to computationally predict CCS values of MS/MS candidate structures and filter those by comparison against the measured CCS value^13^. This process is highly dependent on the accuracy of the analytical IM-MS CCS measurement and the ML prediction errors of CCS values, as isomers often vary by less than 2% in CCS^14^. CCS intra-lab variability is commonly reported to be in the 2% range for DTIM-MS measurements of small molecules when using carefully matched IM-MS parameters. These differences call into question the true efficacy of CCS as a means of filtering similar structure candidates^14^. These challenges are further amplified when attempting to compare CCS data acquired by TWIM-MS or other IM-MS techniques against DTIM-MS databases due to the complexities of the calibration procedures involved^16,17^.

To evaluate the capabilities of these tools when applied in a complex, non-targeted metabolomics scenario, our study centered on the LC-MS metabolomics study of *Caenorhabditis elegans* strains with known mutations to central metabolism pathways. Primary LC-MS analysis was conducted on an Orbitrap platform leveraging its high resolution, robust mass accuracy, and iterative data-dependent acquisition (DDA) capabilities to accurately facilitate elemental formula assignment and provide deep MS/MS coverage. The Orbitrap data set was used to identify differential features of interest to be annotated and generate candidate structures through MS/MS analysis in SIRIUS 4^8^. Following this primary analysis, pooled samples were reanalyzed on a q-IM-TOF instrument to measure CCS values for differential features of interest using LC-TWIM-MS. Candidate structures with ML-predicted CCS values having >3% error against the experimentally measured CCS values were then discarded. Many types of hurdles were encountered through the application of this workflow that, in some cases, precluded effective filtering of candidate structures. Most commonly, the degree of CCS accuracy currently achievable with commercial IM-MS instruments was insufficient for yielding meaningful filtering. In some cases, effective ML CCS predictions were not possible for select adduct species due to limitations in the sizes of the available training sets. Though MS/MS and IM-MS data were successfully collected for 37% of the differential features, fewer than half of these features benefited from some reduction in candidate structures through IM-MS measurements. Despite the attractiveness of leveraging IM-MS CCS measurements to speed up the annotation bottleneck in non-targeted metabolomics, our results point to several areas in need of added research for this approach to reach its full potential.

## EXPERIMENTAL SECTION

### Sample Growth

Two strains of *C. elegans* (*Caenorhabditis* Genetics Center strains VC1265 and RB2347 with alterations to *pyk-1* and *idh-2* respectively) were selected for their mutations to central metabolism pathways and grown in large scale culture plates alongside the PD1074 reference strain samples as previously described^18,19^. Six replicate samples for both test strains were prepared with paired reference samples. Mixed-stage worm populations were harvested, diluted to 200,000 worm aliquots, flash frozen in liquid nitrogen, and stored at −80 °C. Frozen aliquots were then lyophilized with VirTis® BenchTop™ “K” Series Freeze Dryer (SP Industries, Inc.) and stored at −80 °C until extraction.

### Sample Extraction

Samples were evenly divided across two batches to simultaneously accommodate a maximum of 24 samples including controls through the extraction process. Each mutant strain was represented in both batches to account for extraction batch effects. Lyophilized samples were removed from storage at −80 °C and three 2.0 mm zirconium oxide beads and 75 µL volume of 0.5 mm glass beads were added to each sample tube. Samples were homogenized in a Qiagen Tissuelyser II for 3 minutes at 1800 rpm using adapter trays chilled at −80 °C. A sequential extraction was performed starting with the addition of 750 µL LC-MS grade isopropanol (IPA, Fisher Scientific) to each sample. Each sample was lightly vortexed to create a slurry of homogenized samples that was then transferred to a new 2.0 mL centrifuge tube. This process was repeated a second time, so a total volume of 1.5 mL was transferred, leaving behind the homogenizing beads. Samples were vortexed for one minute, and stored overnight (12-15 hours) at −20 °C. Samples were centrifuged at 22100G for 5 minutes and the supernatants transferred to new 2.0 mL centrifuge tubes labeled for reverse phase (RP) chromatography. These extracts were dried in a Labconco CentriVap concentrator until completely dry (4-5 hours) and stored at −80 °C until LC-MS analysis. The remaining pellet was subjected to a second round of extraction using 1.5 mL 80:20 LC-MS grade methanol:water (Fisher Scientific). The pellet and methanol:water mixture was shaken at room temperature (23.0 °C) at 1500 rpm for 30 minutes using a Fisher Scientific Isotemp High Speed Shaker. Samples were again centrifuged at 22100G for 5 minutes and supernatants transferred to new 2.0 mL centrifuge tubes labeled for hydrophilic interaction liquid chromatography (HILIC), dried, and stored at −80 °C.

### Sample Reconstitution and Pooling

Dried IPA extracts were reconstituted with 75 µL IPA containing isotopically labeled lipid standards, vortexed for one minute, centrifuged at 22100G for 5 minutes, and transferred to 300 µL LC-MS vials for RP LC-MS analysis. Methanol:water extracts followed the same reconstitution steps but were instead reconstituted in 80:20 LC-MS grade methanol:water containing isotopically labeled small molecule standards for HILIC LC-MS analysis. Standards used for each extract are described in Table S1. A total of 5 µL from each PD1074 reference sample in the first batch was taken to create a pooled PD1074 sample. Similarly, 5 µL from each mutant strain sample was taken to create a pooled mutant sample. Equal volumes from each pool were mixed to create whole batch pool samples. This pooling process was repeated for batch two.

### LC-MS analysis

Instrument runs began and ended with instrument controls, whole batch pool samples, mutant pool samples, and PD1074 pools. Individual test samples were randomized throughout the middle of the run with three injections of whole batch pool samples interleaved.

IPA samples were LC-MS analyzed using a Thermo Fisher Vanquish chromatograph and Accucore C30 150 × 2.1mm, 2.6 µm column coupled to a Thermo Fisher Orbitrap ID-X Tribrid mass spectrometer. Elution was performed using 40:60 water:acetonitrile with 10 mM ammonium acetate (mobile phase A) and 10:90 acetonitrile:isopropyl alcohol with 10 mM ammonium (mobile phase B). Methanol:water extracts were analyzed on the same system using a Water BEH Acquity UPLC BEH Amide column (2.1×150 mm, 1.7 µm particle size). Elution was performed using 80:20 water:MeCN with 10 mM ammonium formate and 0.1% formic acid (mobile phase A) and MeCN and 0.1% formic acid (mobile phase B). Batches were run in positive, then negative ion polarity before running subsequent batches. Complete chromatographic settings and mass spectrometer parameters for both chromatographic methods are outlined in Tables S2-4.

Full scan MS1 data for each sample was obtained at a resolution setting of 240,000 full-width half maximum (FWHM). MS/MS analysis was performed on whole batch pooled samples using three rounds of iterative DDA (ThermoFisher AcquireX) at a resolution of 30,000 FWH using a 0.8 Da isolation window and stepped HCD collision energies of 15 V, 30 V, and 45 V.

Whole batch pooled samples were re-analyzed on a Waters Synapt G2-S paired to a Water Acquity I-Class UPLC system using matched chromatography to obtain IM drift times from representative samples. Extensive tuning of IMS parameters was performed to limit fragmentation of small molecule species. Briefly, wave voltages and nitrogen gas pressures within the mobility cell were reduced and a wave velocity gradient was implemented to maintain suitable IM resolution. Instrument parameters are described in Table S4. CCS calibration was accomplished using poly-DL-(alanine) (n=2-14) (MilliporeSigma). Multiple reference values for poly-DL-(alanine) were evaluated for their ability to yield accurate CCS values for a wide range of small molecule standards.

### Data Processing

Compound Discoverer 3.1 (ThermoFisher) was used to extract spectral features from the Orbitrap ID-X data sets. Processing steps included retention time (RT) alignment, peak picking, feature grouping, peak integration, and gap filling. The first stage of feature annotation was performed through Compound Discoverer 3.1 using mzCloud (ThermoFisher) and in-house mzVault libraries. For analysis of LCIM-MS data sets acquired on the Waters Synapt G2-S platform, data processing was performed using Progenesis QI 2.4 (Nonlinear Dynamics) to extract features from the raw data and assign CCS values. To match the CCS values of features measured on the Synapt G2-S platform to features measured on the Orbitrap ID-X platform, the RT of internal standards were used to align the chromatographic scales (Figure S1). Following RT correction of the Synapt G2-S data using a linear calibration curve, a 10 ppm *m/z* tolerance and 0.2-minute RT tolerance was used for matching features between platforms manually. In rare cases where multiple features fell within the tolerance window, the Orbitrap ID-X data was manually reviewed to determine elution order to properly match CCS values.

To control for sample variation, a ranked ANOVA approach was used to determine statistically significant differential features for each mutant strain (see Figure S2 and additional text in SI). Features were ranked from most intense to least intense within each sample and binned across 1,000 bins. ANOVA was then performed on the bin number between mutant and control samples using the Galaxy web platform^20^. Bin numbers for features with p<0.05 were used to construct orthogonalized partial least squares discriminant analysis (oPLS-DA) models within PLS_toolbox 8.9.1 (Eigenvector Research, Inc.) with the parameters described in Tables S5-6. Features with top VIP scores were reviewed for proper chromatographic integration and further interrogated for structural identification. At least ten features from each polarity and chromatography mode were investigated for both mutants

### Prediction of Candidate Structures and CCS values

MS/MS data was exported from Compound Discoverer 3.1 in .mgf file format and imported into SIRIUS 4.9.3^8^ to predict the most likely elemental formula and adduct species for each feature based on *m/z*, isotope intensity, and fragmentation pattern. CSI:FingerID^7^ was used within SIRIUS 4.9.3^8^ to generate candidate structures for each feature. Resulting candidate structures were written to .csv files containing the corresponding InChI codes.

To predict CCS values, two data sets were created using the Unified CCS Compendium^11^, one containing all entries for [M+H]^+^ adduct species and another set for all [M-H]^-^ entries (n=644 and 582, respectively). These data sets were then randomly split 75% into training sets and 25% into test sets which were used to construct a support vector regression (SVR)-based ML model using CCSP 2.0^21^ as illustrated in Figure 1. Using the optimized SVR models, CCS values for any given candidate structure could be predicted for [M+H]^+^ and [M-H]^-^ species from their neutral InChI code. The predicted CCS value for each candidate structure was compared against the experimental value obtained from LC-IM-MS data. The accuracy of the predictions was evaluated using the following Equation 1:

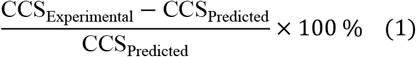

Structures with >3% error were discarded.

**Figure 1.**
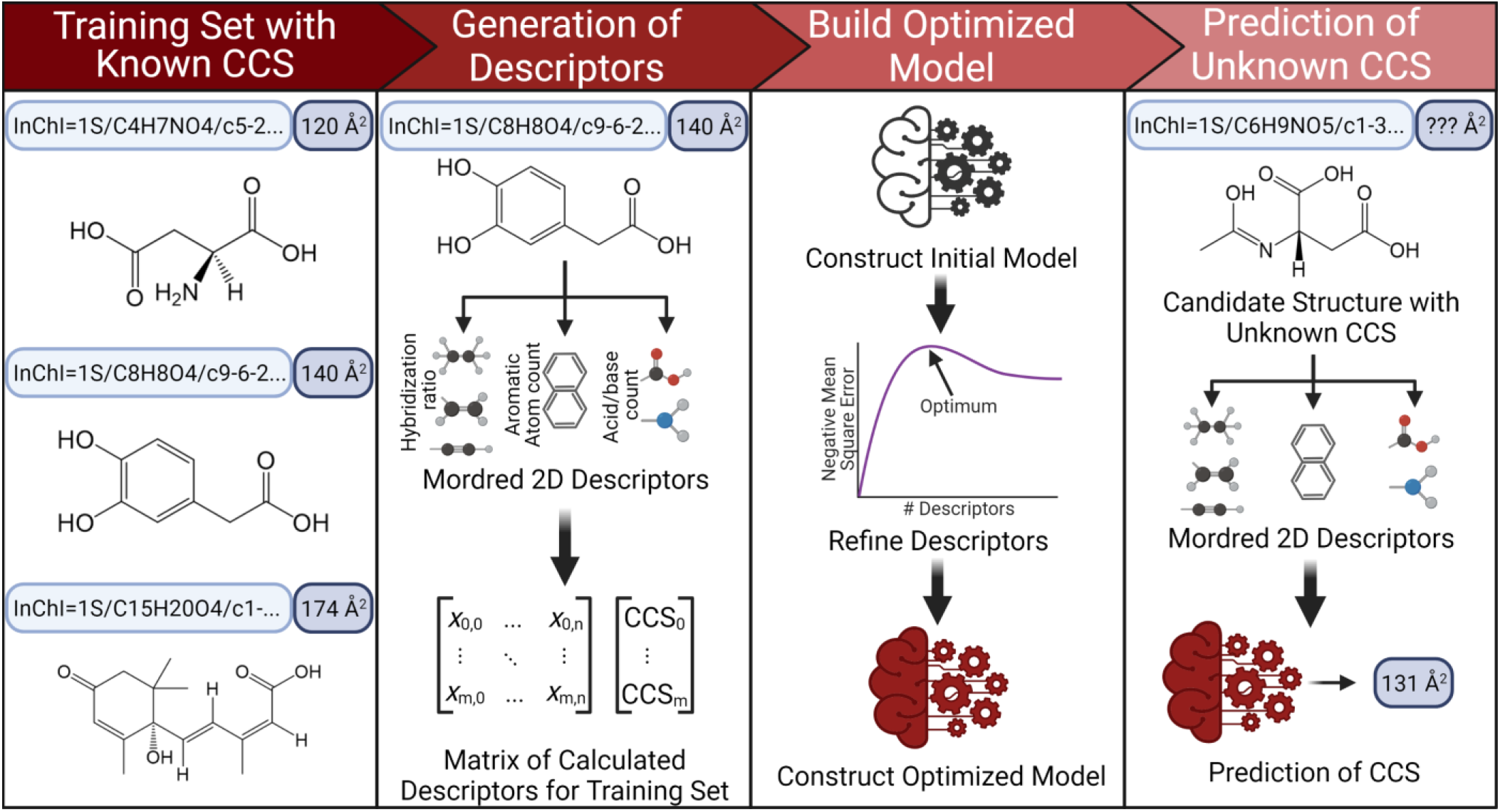
CCSP 2.0 is an SVR-based machine learning algorithm for predicting CCS values from 2D structures represented by InChI strings. Two-dimensional structural data is first used to calculate up to 1613 quantifiable descriptors for each string in the training set through the Python Mordred package, generating a matrix of descriptors matching the matrix of experimental CCS values. Descriptors not applicable to molecules in the training set are culled and an initial SVR model is constructed. Optimization of the model is performed by limiting the number of descriptors used to those with the highest weights, then identifying the fewest descriptors that can be used to yield the lowest CCS prediction error. The calculated descriptors for any candidate 2D structure arising from SIRIUS 4 can then be plugged into the model to yield a predicted CCS value.

## RESULTS AND DISCUSSION

### Challenges Integrating IM-MS in an LC-MS *C. elegans* Metabolomics Pipeline

To evaluate the utility of IM-MS for unknown feature annotation under practical conditions, we developed and tested an experimental workflow that included several innovative steps (Figure 2). Two *C. elegans* strains with mutations to central metabolism pathways (RB2347 and VC1265) were grown alongside reference strain samples in serve as a representative case/control case study for non-targeted metabolomic analysis. The samples consisted of a complex matrix derived from the nematodes, growth media and buffers used. Six samples of the RB2347 and VC1265 mutant strains were prepared together with a growth-matched reference strain sample for each. These samples were sequentially extracted to produce polar and nonpolar fractions. LC-MS analysis of each extract was first performed on an Orbitrap ID-X (ThermoFisher) in both ion polarities. RP chromatography was used for nonpolar extracts and HILIC for polar extracts. This procedure yielded four data sets (RP/HILIC, positive/negative, several Gb each). Orbitrap LC-MS results showed high mass accuracy (<3ppm), whereas iterative DDA AcquireX runs on pooled samples allowed deep MS/MS coverage. From the collected datasets, pre-selection of statistically significant differential features between controls and mutants was performed using a ranked ANOVA approach. Once feature selection *via* ANOVA was completed, it was followed by oPLS-DA modeling. From these oPLS-DA models, a short list of features with the highest VIP scores was created. These features were used as test cases for attempting metabolite annotation using the workflow in Figure 2.A total of 95 features were investigated for structural annotation, 56 of which had MS/MS data collected by iterative DDA. The remaining features were not selected after three rounds of iterative DDA and were not pursued further. Between two and 784 candidate structures were generated for each one of these features in SIRIUS 4^8^ using CSI:FingerID^7^, showing the wide diversity of structures possible from a given MS/MS spectrum when only a few informative fragment ions are present. The CCS for each candidate structure was predicted using the ML CCSP 2.0 algorithm. CCS prediction was not possible for all features as only [M+H]^+^, [M+Na]^+^, and [M-H]^-^ adduct species had sufficient training data available in the Unified CCS Compendium^11^ to build accurate ML models. A total of 66% of the features were detected as one of these adducts, meaning that CCS prediction was not possible for over a third of the features investigated. This was one of the main hurdles faced during this project, as noted in Table S7.

**Figure 2.**
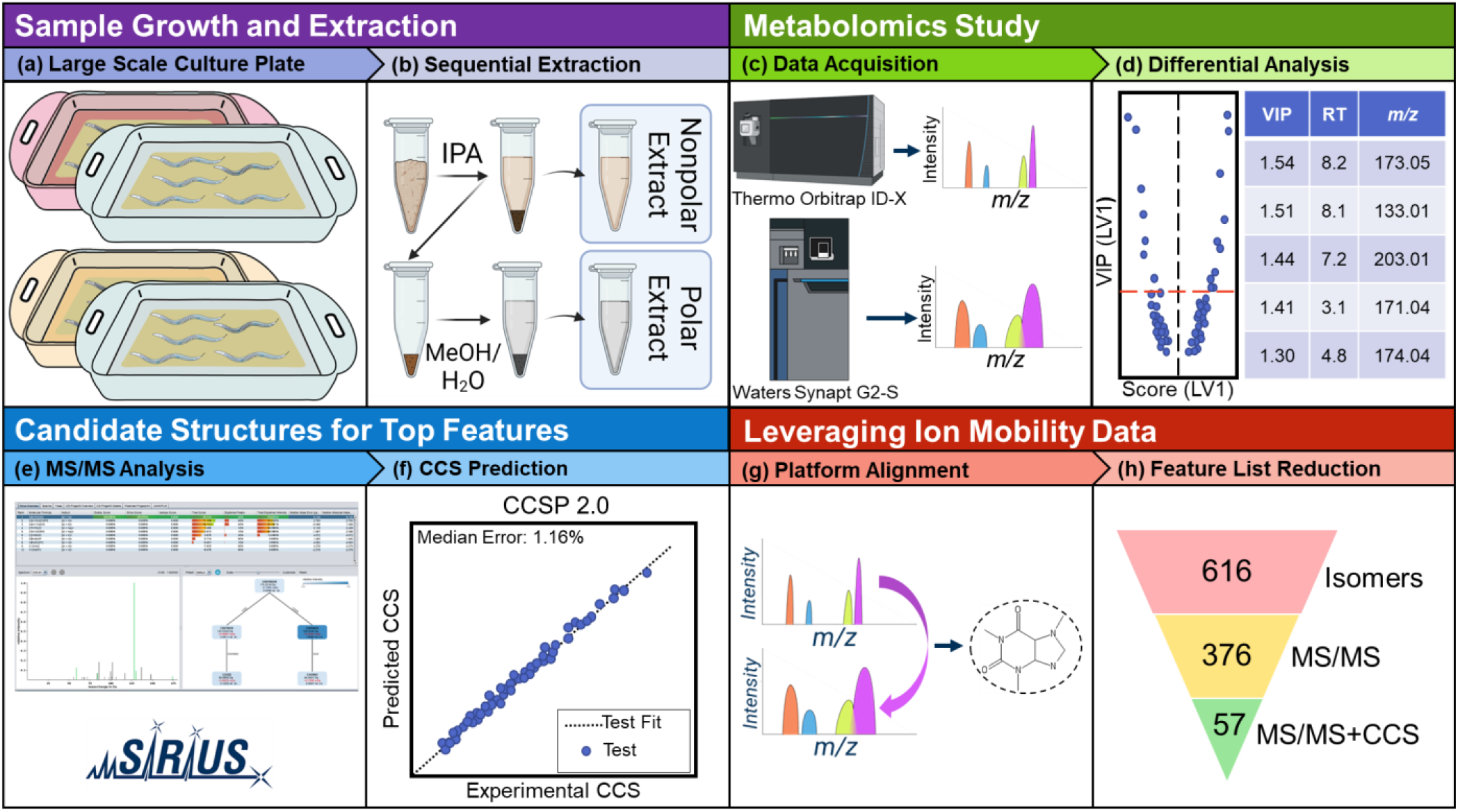
Graphical overview of the metabolomics workflow employed in this study. *C. elegans* mutant strains were grown in tandem with growth-matched PD1074 reference strain samples (a). These cultures were harvested to aliquots of 200,000 worms, lyophilized and then sequentially extracted, yielding nonpolar and polar fractions (b). All samples were analyzed on a Thermo Orbitrap ID-X platform. Pooled samples were re-analyzed on a Waters Synapt G2-S to provide complementary ion mobility data (c). Orbitrap data was used for differential analysis to select significant features of interest (d). Orbitrap MS/MS data was imported to SIRIUS 4^*8*^ to generate candidate structures using CSI:FingerID^*7*^ (e). The CCS value for each candidate structure produced by SIRIUS 4 was predicted using CCSP 2.0, an SVR-based machine learning algorithm trained with the McLean Unified CCS Compendium^*11*^ data (f). Synapt ion mobility data was matched to Orbitrap data to provide experimental CCS values for differential features (g) which were then used to eliminate candidate structures falling outside the 3% error band (h).

Pooled *C.elegans* samples were analyzed on the Synapt G2S platform to collect TWIM-MS data and provide experimentally measured CCS values for features of interest. Matched chromatography was used, though RT correction using internal standards was required. Features detected in the Orbitrap ID-X platform could then be aligned to those measured on the Synapt G2-S by matching them with 10 ppm and 0.2 min. RT error after correction. A total of 55 out of the 95 differential features were matched using these criteria and could be assigned experimental CCS values. The lowest observed success rate was for lipid species in the RP dataset. Some of these were only observed as dimeric ions (*e.g*. [2M+H]^+^) on the Orbitrap ID-X but not detected on the Synapt G2-S. It is possible that these dimeric ions either do not form as readily in the Synapt instrument ion source, or that these species do not survive more complex ion optics, which include the higher-pressure IM cell, without fragmenting. Features from the HILIC Orbitrap dataset were also sometimes missing in the Synapt data set, likely due to unwanted ion activation in the mobility stage or differences in ion transmission between platforms.

Fragmentation products observed for labile metabolites during LC-TWIM-MS method development with standard metabolites included amino acid deamination, decarboxylation of organic acids, and fragmentation of compounds such as hippuric acid. Extensive tuning of the IM parameters was performed to minimize unwanted fragmentation of small molecules occurring during ion transport from the higher vacuum quadrupole region into the higher-pressure mobility cell of the instrument. The final IM-MS parameters chosen to minimize fragmentation are given in Table S4. Another reason that could lead to the lack of matched features between the Orbitrap and Synapt systems is related to differences in their mass resolution. It is possible that spectral overlaps in the lower resolution Synapt platform resulted in the mass errors that caused features to fall outside the 10 ppm window.

Candidate structures for the investigated differential features could be filtered once these were assigned experimental CCS values from TWIM-MS data, candidate structures from Orbitrap MS/MS data, and ML-predicted CCS values. The differences observed between ML-predicted *vs*. experimental CCS values is a complex convolution of several factors that include: 1) the experimental TWIM-MS CCS measurement and calibration errors and 2) the SVR model errors inherent to ML predictions. Regarding the latter, multiple attempts to accurately predict the uncertainty of SVR models *via* bootstrap approaches^22–24^ have been attempted, but none have yet been implemented in the context of CCS prediction algorithms. The considerable number of molecular descriptor inputs and the lack of comprehensive training data complicate the generation of point-wise confidence intervals for each metabolite of interest. Consequently, CCS prediction algorithms typically rely on summary statistics such as the root mean square error (RMSE) and the median relative error (MRE) to set blanket exclusion thresholds. Most CCS prediction algorithms in the literature suggest a 3% threshold for matching the experimentally measured CCS and the predicted CCS values of their test sets^25–27^. Though more restrictive (1%) and more permissive^13^ (4%) thresholds have also been investigated, a 3% filtering threshold was adopted here based on the calibration errors observed in our experiments and to allow comparison between our results and those previously published.

### Machine Learning CCS Predictions

CCSP 2.0, our inhouse ML algorithm for CCS prediction, encodes molecular structure through InChI strings and uses molecular descriptors for such structures to generate SVR models that accurately predict CCS for unknowns (Figure 1). This ML tool was created in a Jupyter notebook with freely available open-source Python packages for maximum accessibility and to enable the user to modify the code as needed. Prior to applying CCSP 2.0 to the *C. elegans* IM-MS dataset, its prediction accuracy was estimated through an external validation procedure. The CCS prediction error was 1.14% for [M-H]^-^ ions and 1.56% for [M+H]^+^ ions. These errors corresponded to RMSE values of 4.268 Å^2^ and 5.694 Å^2^, respectively, which were comparable to or better than other published CCS prediction methods^12,13^. Further CCS prediction cross validation results are provided in Figure S3, showing an excellent correlation between measured and predicted CCS values. This level of prediction accuracy, together with the flexible CCSP 2.0 code environment allows other research teams to adapt the training set to their specific applications and to make use of hybrid calibration strategies combining experimental and predicted CCS values, as discussed in the next section and in Figure 3.

**Figure 3.**
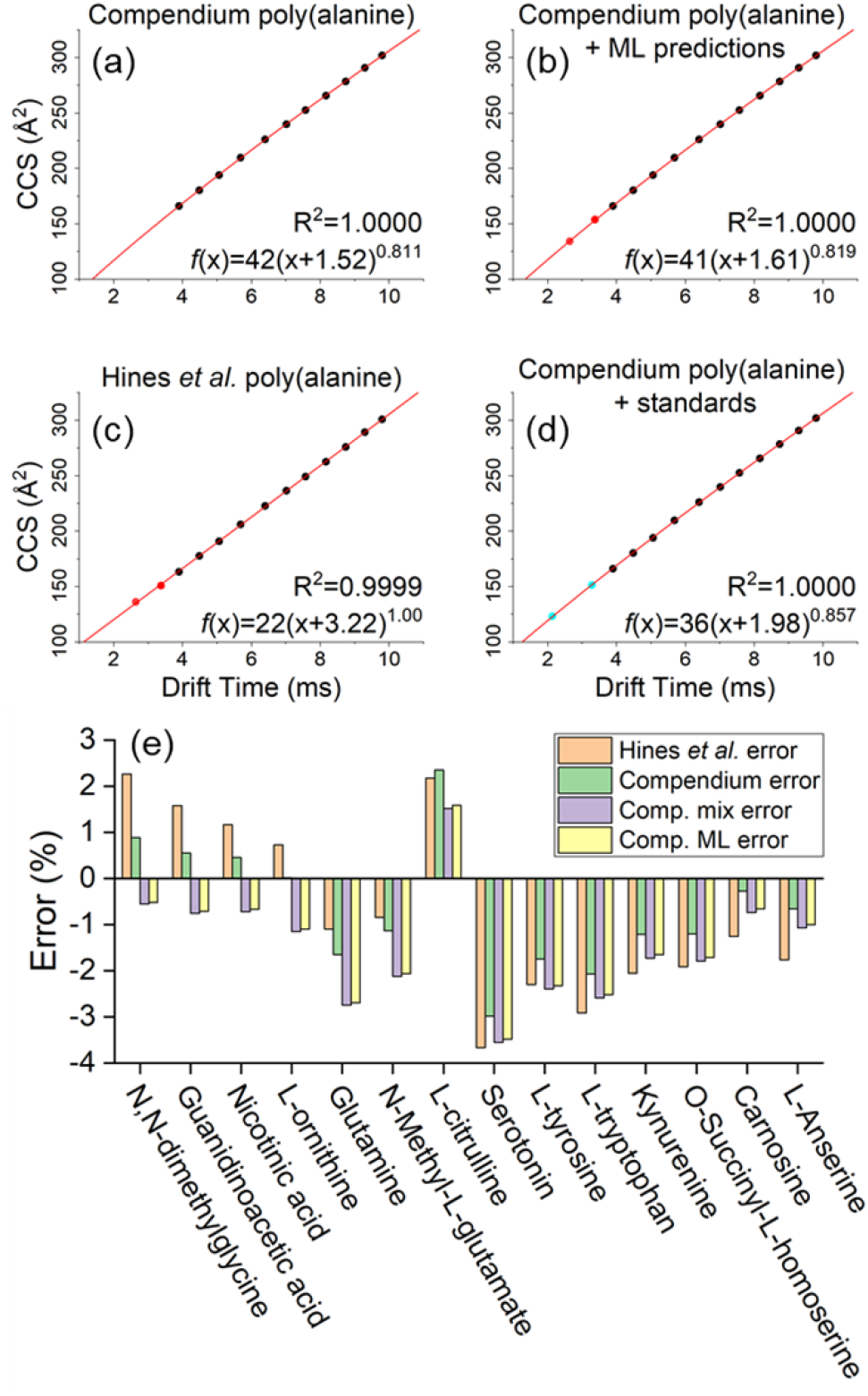
The accuracy of four CCS calibration strategies were evaluated using a panel of 14 standards covering the 104-241 *m/z* range. Plot (a) shows a power law calibration curve using Unified CCS Compendium^11^ data covering poly(alanine) oligomer lengths *n*=4-14 (black points). To further extend the range of calibration, this series may be extended using ML predicted CCS values for poly(alanine) *n*=2-3 (red points) as shown in plot (b) or by using poly(alanine) data from an alternate source (c). Alternatively, the poly(alanine) series may be supplemented with additional standards as in (d) where dimethylglycine and tryptophan (blue points) were used. CCS values for these 14 standards were calculated using each of the four power-law calibration curves, and then compared against Unified CCS Compendium^11^ reported values for each standard. Errors for each calibration procedure are shown in (e).

### Optimizing TWIM Calibration Accuracy for Diverse Da-tasets

Minimizing CCS calibration errors is a critical component in obtaining the most accurate CCS values for filtering structures generated in SIRIUS 4. TWIM-MS CCS measurements (^TW^CCSN228). require calibration of drift times with a series of compounds with known CCS values, typically measured by DTIM-MS (^DT^CCSN2). Poly-DL-(alanine) oligomers are commonly used as a calibrant mixture because they exhibit relatively little chemical class bias. This makes this calibrant well suited for non-targeted IM-MS analysis^29–32^. The source of ^DT^CCSN2 reference values plays a large role in the quality of the results produced by TWIM-MS following drift time calibration. Ideally, calibration should be performed using data from the database source that will ultimately be used for comparisons. For example, TWIM-MS drift times calibrated using Unified CCS Compendium^11 DT^CCSN2 values will have the highest agreement with other CCS data from this source if compared to data from alternative sources such as AllCCS^13^ and CCSbase^12^. However, the CCS reference database of choice often may not include complete data for the desired calibrant series. In our case, the Unified CCS Compendium^11^ only reports CCS values for poly(alanine) down to the *n*=4 oligomer ([M+H]^+^ = *m/z* 303.1668) though the range of differential metabolites investigated spanned *m/z* values between 70 and 1050. While powerlaw calibrations can be used to extrapolate CCS values outside of the calibrated range, it is possible that including additional calibration points to cover the expected *m/z* range may further improve results. To achieve greater CCS calibration coverage, we tested four different calibration methods. The basic method only used the poly(alanine) CCS values from the Unified CCS Compendium^11^. A second method supplemented the Unified CCS Compendium poly(alanine) data with Compendium values from tryptophan and dimethylglycine. A third calibration method used the full poly(alanine) CCS reference data from Hines *et al*.^30^ which includes the *n*=2-3 oligomers. The fourth and last calibration approach used ML-predicted CCS values for the *n*=2-3 oligomers obtained when using Unified CCS Compendium^11^ data for the training set.

The accuracy of these four calibration methods was compared by measuring CCS values of 14 chemical standards covering the *m/z* 104-241 range in positive ion polarity. A single IM-MS data set was acquired and then calibrated using each of the four calibration methods. Calculated CCS values were then compared against those reported in the Unified CCS Compendium. Resulting errors are shown in Figure 3. On average, the poorest agreement (% average absolute error=1.84) was observed when calibrating with data from Hines *et al*. and comparing against the Unified CCS Compendium (Figure 3c). These differences likely arise from benchmarking data calibrated with reference values from one laboratory against the values from another laboratory. It may also arise from the inherently different TWIM and DTIM gas-phase separation mechanisms. CCS data calibrated using only CCS Compendium poly(alanine) values yielded the lowest average absolute error (1.23%, Figure 3a). Calibration with additional standards (Figure 3d) produced an absolute error of 1.68% on average, and calibration supplemented with ML-predicted points performed similarly to the approach involving additional standards, with an average absolute error of 1.62% (Figure 3b). The latter results illustrate the possibility of enhancing TWIM calibrations with ML-derived values in cases where the reference database is incomplete. Figure 3e shows the results for each of the individual standard compounds tested. Based on these results, the method depicted in Figure 3a was selected moving forward as it yielded absolute errors ranging from 0.02% to 2.98%. Based on these calibration errors, the chosen CCS threshold of 3% for comparing predicted and experimental CCS values was in line with literature results.

### Distribution of ML CCS Prediction Errors for SIRIUS 4-derived Structures

The errors of the experimentally measured *vs*. ML predicted CCS values for each candidate structure generated in SIRIUS 4 were calculated for all 19 differential features investigated. Histograms of these measured *vs*. predicted CCS error distributions are shown in Figure 4. The overall success of the annotation workflow CCS filtering step can be evaluated from the results shown in Figure 4a. A total of 1580 candidate structures were filtered out from the original 3495 structures proposed by SIRIUS 4. Interestingly, an overall shift towards negative errors was observed in the histogram. This effect can likely be attributed to the choice of polyalanine for converting TWIM drift times to CCS values (Figure 3e). Bias in the calibration may be introduced by the nuanced interactions of the chosen calibrants with the transient electric fields present in the TWIM drift cell. These interactions change based on the chemical class in ways not accounted for in typical power-law drift time calibrations, such as the one used here^16^. As a result, chemical class differences between calibrant and analyte manifest as CCS biases in the same direction, positive or negative^17^. While poly(alanine) serves as a general calibrant offering reasonably good structural similarity for a variety of compounds^29–32^, other calibrants could likely exhibit better calibrant-analyte structural similarity. This is important for non-targeted metabolomics studies dealing with a variety of chemical classes, such as those involved in central metabolism mutant *C. elegans* strains. However, chemical class matching of the calibrants to the unknowns is only possible with *a priori* knowledge of the chemical class of the investigated features. Moreover, CCS databases often lack sufficient representation from compounds of a certain class of interest, such as adduct ions that could match the ionic species detected.

**Figure 4.**
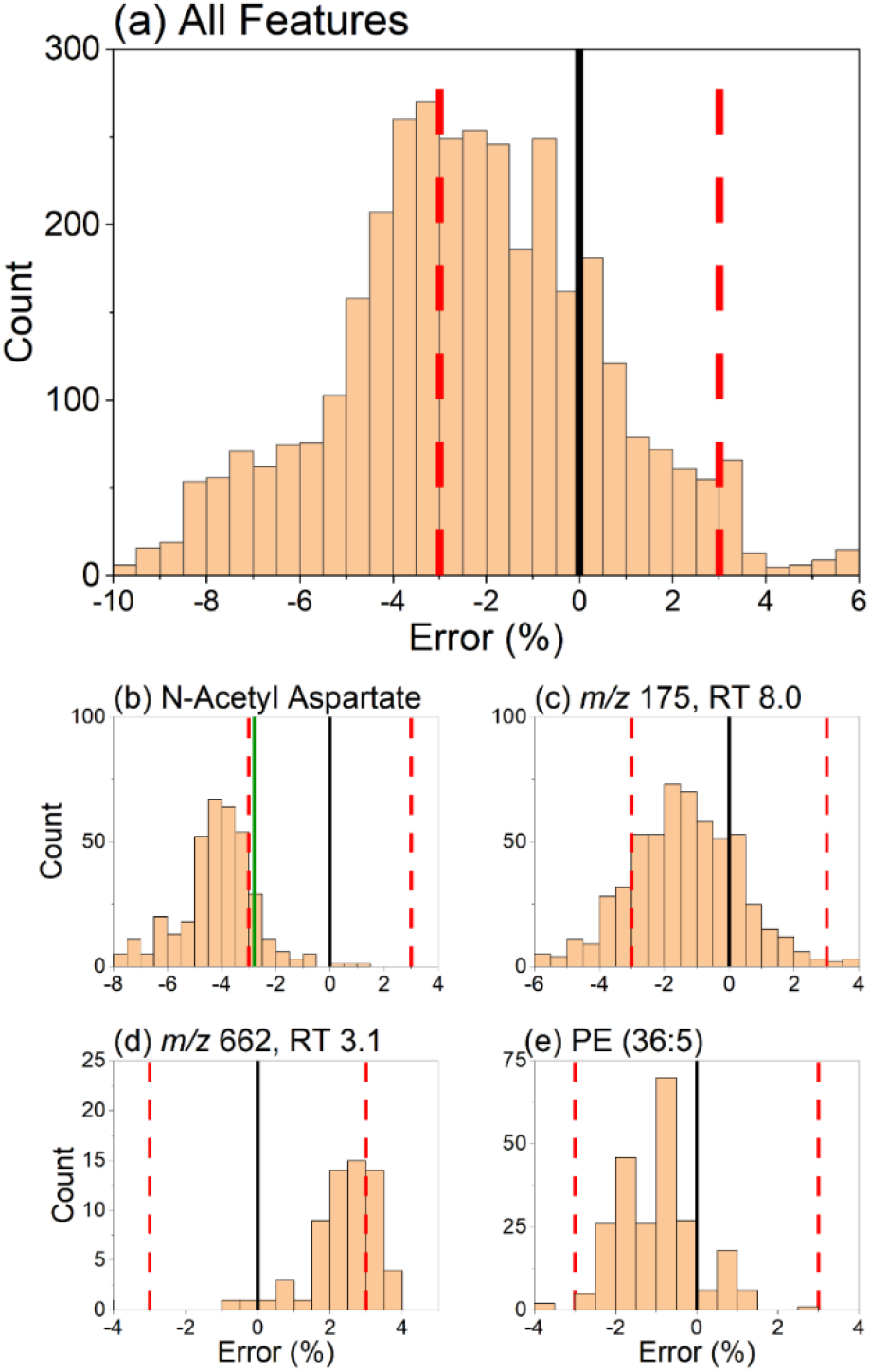
Histogram describing the percent error between ML-predicted CCS values of SIRIUS 4 candidate structures (*n*=3495) *vs*. experimentally measured CCS. The error in these predictions was defined as in Equation 1. Shown in (a) is the summation of errors for all the 19 differential features investigated, demonstrating an overall filtering success of 45.2% (1580 out of 3495). A negative bias towards lower-than-predicted experimental CCS values was observed. Individual errors for four features of interest are shown in plots (b) through (e). The solid black line represents the experimentally measured CCS value. The red dashed lines represent the ±3% CCS cutoff. The solid green line represents the error of the correct compound validated with a chemical standard.

Error histograms for individual features displayed similar Gaussian-like distributions due to the high similarity of the structures generated from *in silico* analysis of a given MS/MS spectrum. Many candidate structures fell within ±3% of the center of each error distribution, and thus the percent of features filtered by CCS was highly dependent on how far the center of such error distribution was shifted from the experimental CCS value. A best-case scenario is shown in Figure 4b where the distribution was centered at roughly −4% error. In this case, the correct structural annotation (N-acetyl aspartate) fell on the right shoulder of the distribution, just within the ±3% CCS cutoff. The identity of the correct compound was confirmed with a pure standard and through searches in mzCloud during data processing. N-acetyl aspartate represents an almost ideal case where CCS can filter >50% of candidate structures while retaining the correct annotation. Out of the 57 candidate structures within the ±3% CCS cutoff (Figure 4b), N-acetyl aspartate was the #1 ranked SIRIUS 4 candidate. This example, however, is not typical. A more frequent scenario is shown in Figure 4c for a yet-unidentified feature at *m/z* 175.0713 and RT 8.02 minutes, where only 19% of candidate structures fell outside the ±3% CCS cutoff. Figure 4d depicts a case of an unknown with a CCS error distribution shifted towards positive values. In this case, 33% of candidate structures were filtered. The MS/MS and CCS data on the remaining 45 structures were insufficient to make a conclusive annotation. These results highlight the importance of reducing the CCS cutoff value by improving overall CCS measurement accuracy.

Lipid species are especially challenging due to the multitude of possible isomeric structures of similar molecular size and shape. *In silico* prediction of structures from MS/MS spectra yields fewer structures than for other classes of compounds, but these all have very similar predicted CCS values. This results in a narrower error distribution that is more difficult to filter by TWIM-MS measurements.

### Structural Annotation of Differential Features in *C.elegans* Mutants

A total of 95 features were identified as differential between mutant and control strains and further investigated with the proposed workflow. Matched MS/MS and CCS data were successfully collected for 34 of these features. Of these 34, only 19 could be processed through the entire workflow. The effectiveness of CCS filtering for each of these 19 features is shown in Figure 5. Both the number of candidate structures generated and the percent of structures filtered by CCS varied substantially on a case-to-case basis. An average of 183.9 candidate structures were generated for each feature with a mean reduction of 28% of structures on an individual basis using CCS filtering. Notably, six features had no reduction in candidate structures, meaning all structures had predicted CCS values within 3% of the measured CCS. For a differential feature at *m/z* 738 and RT 3.1 min in the RP data set, only two of the 233 candidate structures could be filtered. This shows that while many structures may be possible for a given formula, MS/MS data will often constrain the possibilities to those of highly similar molecular size and shape, and therefore CCS values. For the 15 features which could not be processed through the full annotation workflow, several unique failure points were identified. As previously mentioned, sufficient training data was not available in many cases to construct accurate ML models for the measured type of ion adduct. Other points of failure include: 1) no candidate structures from MS/MS data produced by SIRIUS 4, 2) TWIM-MS ion rollover identified as an abnormally small CCS value, and 3) incorrect processing of TWIM-MS drift time data by Progenesis software yielding incorrect drift times. A full description of identified failure points for each of the 15 failed features is given in Table S7. Though ion rollover and incorrect arrival time assignment could be rectified through follow-up experiments and manual data processing, it is important to note the difficulty of designing an automated workflow that would work universally on every feature without exception.

**Figure 5.**
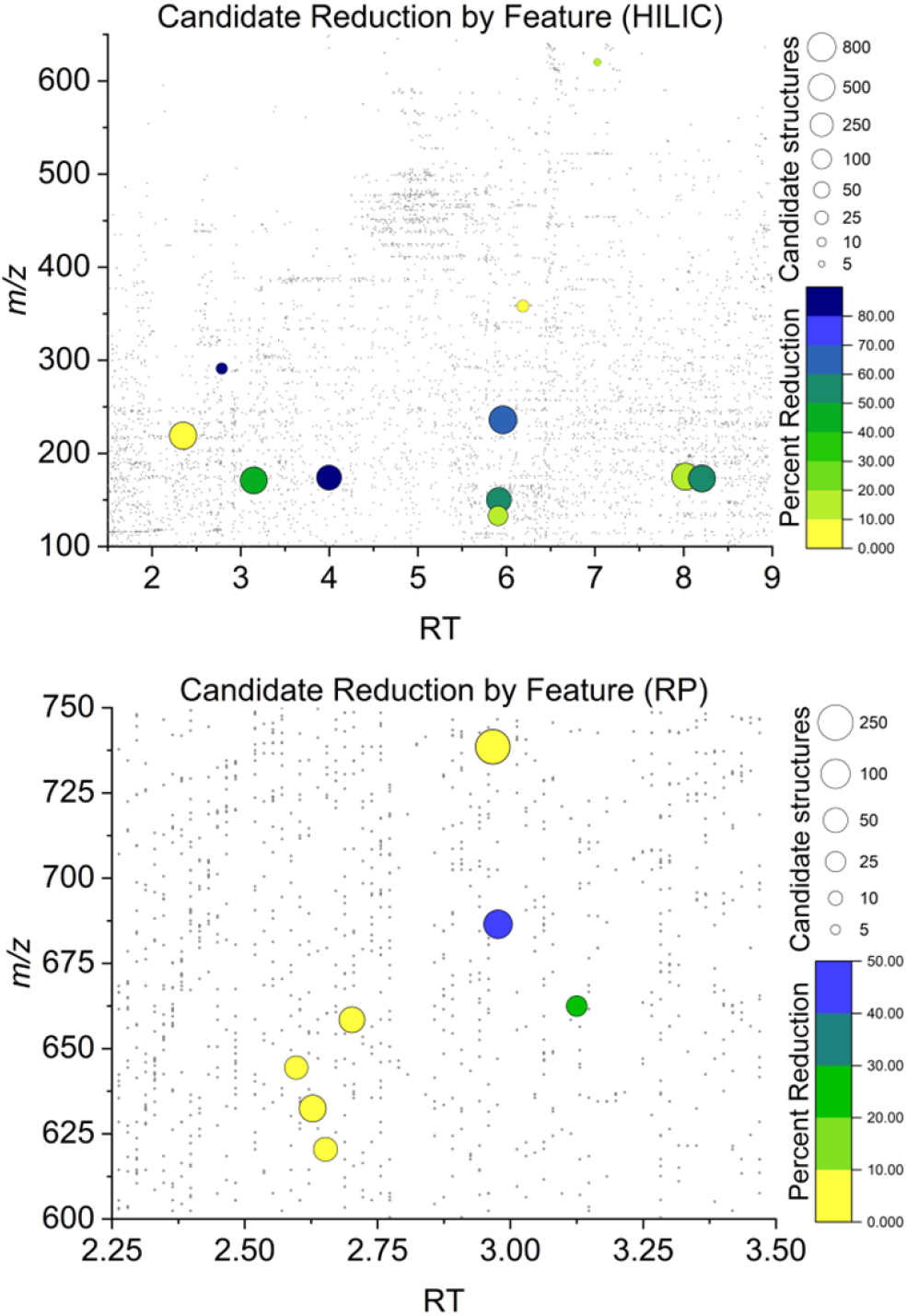
Overall results from CCS assisted annotation of unknown features. Each investigated feature is represented by a bubble with the size defined by the total number of MS^2^ generated candidate structures, and color defined by the percent of total candidates which could be discarded by their CCS value. A background of gray dots represents all detected features.

Certain assignments were confirmed for three of the 19 successfully investigated features using pure analytical standards by matching RT and drift time from the Synapt G2-S and matching MS/MS spectra from the Orbitrap ID-X. Two of these features were also identified first within mzCloud: N-acetyl aspartate and guanine. While these compounds do not necessarily benefit from CCS candidate structure filtering as putative identifications can be assigned by database MS/MS matching, they do serve as effective validations of the workflow (Figure S4). The MS/MS spectra from N-acetyl aspartate generated 376 candidate structures, 85% of which were filtered by CCS. Guanine produced 365 candidate structures, 54% of which were filtered by CCS. The measured CCS values for both features were within 3% of the predicted CCS for the correct structure.

Perhaps the most useful application of the proposed workflow was for the feature at *m/z* 175 and RT 8.3 where only two candidate structures were generated. One of these two structures could be discarded by CCS, leaving only the possibility for allantoic acid, which was later confirmed by *m/z*, RT and MS/MS matching to a pure standard. This was the only such case where the application of the workflow resulted in only a single remaining candidate from numerous possibilities in the starting set. More commonly, dozens to hundreds of candidate structures remained following CCS filtering; still too many to attempt matching to a chemical standard, highlighting the need for the application of additional techniques such as NMR spectroscopy to finalize the annotation process.

## CONCLUSIONS

In this manuscript we have explored the utility of augmenting non-targeted metabolomics workflows with an ion mobility candidate structure filtering step to establish the performance of this approach under “real world” conditions involving complex samples. The results indicated that the number of filtered structures was highly variable and limited mainly by the accuracy of CCS measurements and ML predictions achievable with current instrumentation and calibration approaches. With the current CCS cutoff of 3% typically used in the literature, filtering eff0069cacy depended on chemical class and the amount of information that could be gleaned from MS/MS fragment ions. On average, a mean reduction of 28% of structures on an individual basis was observed.

As higher resolution and higher accuracy IM-MS instrumentation is developed, and CCS prediction ML models for additional ionic species become available, it can be expected that the utility of CCS candidate filtering for metabolite identification will also improve. While many recent advancements have resulted in IM-MS platforms with considerably higher resolution, our work highlights the need for improvements in CCS accuracy, not only resolving power. A greater agreement in measured CCS between platforms is also highly desirable. New IM-MS platforms, such as trapped IM, cyclic IM and structures for lossless ion manipulations (SLIM) IM, offer much improved separation capabilities compared to the more standard TWIM-MS platform tested here, but the variance between DTIM CCS values and those platforms is still >1% in most cases. Our results highlight that the current level of accuracy of CCS values is still insufficient for reaching the full potential of IM-MS based metabolite annotation efforts. Improvements in ML prediction of CCS values through better training sets will also likely reduce one of the main sources of variance in the CCS filtering approach.

## ASSOCIATED CONTENT

## Supporting Information

Chromatographic matching curves, data processing steps for LC-MS data, performance metrics for CCS predictions, individual examples of CCS filtering performance, list of internal standards used, details of chromatographic gradients, MS instrument parameters, oPLS-DA parameters, genetic algorithm parameters, list of failure points observed for each investigated metabolite failing to complete the annotation workflow.

The Supporting Information is available free of charge on the ACS Publications website.

## AUTHOR INFORMATION

### Author Contributions

CKA performed all LC-MS and IM-MS experiments, BMG provided help with LC-MS experiments and extraction protocols, MR and CKA conducted CCS predictions. GJG and AOS grew *C.elegans* samples and maintained colonies. AMM and LMM conducted data processing steps and statistical analysis. FEL, ASE and FMF provided overall guidance in experimental design and research strategy. CKA and FMF wrote the manuscript and the rest of the authors edited it. All authors have given approval to the final version of the manuscript.

## ACKNOWLEDGMENT

Research reported in this manuscript was supported by the National Institutes of Health Award Number 1U2CES030167-01 (ASE contact PI) and by grant 1R01CA218664-01 to FMF. We thank Kenneth M. Merz Jr. and his team for insightful discussions. Elements of Figures 1, 2 and S2 were created with BioRender.com

## REFERENCES

(1) Wishart, D. S.; Guo, A.; Oler, E.; Wang, F.; Anjum, A.; Peters, H.; Dizon, R.; Sayeeda, Z.; Tian, S.; Lee, B. L.; Berjanskii, M.; Mah, R.; Yamamoto, M.; Jovel, J.; Torres-Calzada, C.; Hiebert-Giesbrecht, M.; Lui, V. W.; Varshavi, D.; Varshavi, D.; Allen, D.; Arndt, D.; Khetarpal, N.; Sivakumaran, A.; Harford, K.; Sanford, S.; Yee, K.; Cao, X.; Budinski, Z.; Liigand, J.; Zhang, L.; Zheng, J.; Mandal, R.; Karu, N.; Dambrova, M.; Schiöth, H. B.; Greiner, R.; Gautam, V. HMDB 5.0: The Human Metabolome Database for 2022. Nucleic Acids Res 2021, 50 (D1), D622–D631. https://doi.org/10.1093/nar/gkab1062.

(2) Sud, M.; Fahy, E.; Cotter, D.; Brown, A.; Dennis, E. A.; Glass, C. K.; Merrill, A. H.; Murphy, R. C.; Raetz, C. R. H.; Russell, D. W.; Subramaniam, S. LMSD: LIPID MAPS Structure Database. Nucleic Acids Res 2007, 35 (Database issue), D527–532. https://doi.org/10.1093/nar/gkl838.

(3) Horai, H.; Arita, M.; Kanaya, S.; Nihei, Y.; Ikeda, T.; Suwa, K.; Ojima, Y.; Tanaka, K.; Tanaka, S.; Aoshima, K.; Oda, Y.; Kakazu, Y.; Kusano, M.; Tohge, T.; Matsuda, F.; Sawada, Y.; Hirai, M. Y.; Nakanishi, H.; Ikeda, K.; Akimoto, N.; Maoka, T.; Takahashi, H.; Ara, T.; Sakurai, N.; Suzuki, H.; Shibata, D.; Neumann, S.; Iida, T.; Tanaka, K.; Funatsu, K.; Matsuura, F.; Soga, T.; Taguchi, R.; Saito, K.; Nishioka, T. MassBank: A Public Repository for Sharing Mass Spectral Data for Life Sciences. J Mass Spectrom 2010, 45 (7), 703–714. https://doi.org/10.1002/jms.1777.

(4) Bowen, B. P.; Northen, T. R. Dealing with the Unknown: Metabolomics and Metabolite Atlases. Journal of the American Society for Mass Spectrometry 2010, 21 (9), 1471–1476. https://doi.org/10.1016/j.jasms.2010.04.003.

(5) Silva, R. R. da; Dorrestein, P. C.; Quinn, R. A. Illuminating the Dark Matter in Metabolomics. PNAS 2015, 112 (41), 12549–12550. https://doi.org/10.1073/pnas.1516878112.

(6) Vaniya, A.; Fiehn, O. Using Fragmentation Trees and Mass Spectral Trees for Identifying Unknown Compounds in Metabolomics. Trends Analyt Chem 2015, 69, 52–61. https://doi.org/10.1016/j.trac.2015.04.002.

(7) Dührkop, K.; Shen, H.; Meusel, M.; Rousu, J.; Böcker, S. Searching Molecular Structure Databases with Tandem Mass Spectra Using CSI:FingerID. PNAS 2015, 112 (41), 12580–12585. https://doi.org/10.1073/pnas.1509788112.

(8) Dührkop, K.; Fleischauer, M.; Ludwig, M.; Aksenov, A. A.; Melnik, A. V.; Meusel, M.; Dorrestein, P. C.; Rousu, J.; Böcker, S. SIRIUS 4: A Rapid Tool for Turning Tandem Mass Spectra into Metabolite Structure Information. Nat Methods 2019, 16 (4), 299–302. https://doi.org/10.1038/s41592-019-0344-8.

(9) Blaženović, I.; Kind, T.; Ji, J.; Fiehn, O. Software Tools and Approaches for Compound Identification of LC-MS/MS Data in Metabolomics. Metabolites 2018, 8 (2), 31. https://doi.org/10.3390/metabo8020031.

(10) Paglia, G.; Astarita, G. Metabolomics and Lipidomics Using Traveling-Wave Ion Mobility Mass Spectrometry. Nature Protocols 2017, 12 (4), 797–813. https://doi.org/10.1038/nprot.2017.013.

(11) Picache, J. A.; Rose, B. S.; Balinski, A.; Leaptrot, K. L.; Sherrod, S. D.; May, J. C.; McLean, J. A. Collision Cross Section Compendium to Annotate and Predict Multi-Omic Compound Identities. Chem Sci 2019, 10 (4), 983–993. https://doi.org/10.1039/c8sc04396e.

(12) Ross, D. H.; Cho, J. H.; Xu, L. Breaking Down Structural Diversity for Comprehensive Prediction of Ion-Neutral Collision Cross Sections. Anal. Chem. 2020, 92 (6), 4548–4557. https://doi.org/10.1021/acs.analchem.9b05772.

(13) Zhou, Z.; Luo, M.; Chen, X.; Yin, Y.; Xiong, X.; Wang, R.; Zhu, Z.-J. Ion Mobility Collision Cross-Section Atlas for Known and Unknown Metabolite Annotation in Untargeted Metabolomics. Nat Commun 2020, 11 (1), 4334. https://doi.org/10.1038/s41467-020-18171-8.

(14) Dodds, J. N.; May, J. C.; McLean, J. A. Investigation of the Complete Suite of the Leucine and Isoleucine Isomers: Towards Prediction of Ion Mobility Separation Capabilities. Anal Chem 2017, 89 (1), 952–959. https://doi.org/10.1021/acs.analchem.6b04171.

(15) Paglia, G.; Williams, J. P.; Menikarachchi, L.; Thompson, J. W.; Tyldesley-Worster, R.; Halldórsson, S.; Rolfsson, O.; Moseley, A.; Grant, D.; Langridge, J.; Palsson, B. O.; Astarita, G. Ion Mobility Derived Collision Cross Sections to Support Metabolomics Applications. Anal Chem 2014, 86 (8), 3985–3993. https://doi.org/10.1021/ac500405x.

(16) Richardson, K.; Langridge, D.; Dixit, S. M.; Ruotolo, B. T. An Improved Calibration Approach for Traveling Wave Ion Mobility Spectrometry: Robust, High-Precision Collision Cross Sections. Anal. Chem. 2021, 93 (7), 3542–3550. https://doi.org/10.1021/acs.anal-chem.0c04948.

(17) Hines, K. M.; May, J. C.; McLean, J. A.; Xu, L. Evaluation of Collision Cross Section Calibrants for Structural Analysis of Lipids by Traveling Wave Ion Mobility-Mass Spectrometry. Anal Chem 2016, 88 (14), 7329–7336. https://doi.org/10.1021/acs.analchem.6b01728.

(18) Shaver, A. O.; Gouveia, G. J.; Kirby, P. S.; Andersen, E. C.; Edison, A. S. Culture and Assay of Large-Scale Mixed-Stage Caenorhabditis Elegans Populations. JoVE (Journal of Visualized Experiments) 2021, No. 171, e61453. https://doi.org/10.3791/61453.

(19) Shaver, A. O.; Garcia, B. M.; Gouveia, G. J.; Morse, A. M.; Liu, Z.; Asef, C. K.; Borges, R. M.; Leach, F. E.; Andersen, E. C.; Amster, I. J.; Fernández, F. M.; Edison, A. S.; McIntyre, L. M. An Anchored Experimental Design and Meta-Analysis Approach to Address Batch Effects in Large-Scale Metabolomics. bioRxiv March 27, 2022, p 2022.03.25.485859. https://doi.org/10.1101/2022.03.25.485859.

(20) Afgan, E.; Baker, D.; Batut, B.; van den Beek, M.; Bouvier, D.; Čech, M.; Chilton, J.; Clements, D.; Coraor, N.; Grüning, B. A.; Guerler, A.; Hillman-Jackson, J.; Hiltemann, S.; Jalili, V.; Rasche, H.; Soranzo, N.; Goecks, J.; Taylor, J.; Nekrutenko, A.; Blankenberg, D. The Galaxy Platform for Accessible, Reproducible and Collaborative Biomedical Analyses: 2018 Update. Nucleic Acids Research 2018, 46 (W1), W537–W544. https://doi.org/10.1093/nar/gky379.

(21) Rainey, M. A.; Watson, C. A.; Asef, C. K.; Foster, M. R.; Baker, E. S.; Fernández, F. M. CCS Predictor 2.0: An Open-Source Jupyter Notebook Tool for Filtering Out False Positives in Metabolomics. bioRxiv August 9, 2022, p 2022.08.09.503345. https://doi.org/10.1101/2022.08.09.503345.

(22) De Brabanter, K.; De Brabanter, J.; Suykens, J. A. K.; De Moor, B. Approximate Confidence and Prediction Intervals for Least Squares Support Vector Regression. IEEE Trans Neural Netw 2011, 22 (1), 110–120. https://doi.org/10.1109/TNN.2010.2087769.

(23) Harrington, P. de B. Automated Support Vector Regression. Journal of Chemometrics 2017, 31 (4), e2867. https://doi.org/10.1002/cem.2867.

(24) Lins, I. D.; Droguett, E. L.; Moura M. das C.; Zio, E.; Jacinto, C. M. Computing Confidence and Prediction Intervals of Industrial Equipment Degradation by Bootstrapped Support Vector Regression. Reliability Engineering & System Safety 2015, 137, 120–128. https://doi.org/10.1016/j.ress.2015.01.007.

(25) Zhou, Z.; Shen, X.; Tu, J.; Zhu, Z.-J. Large-Scale Prediction of Collision Cross-Section Values for Metabolites in Ion Mobility-Mass Spectrometry. Anal Chem 2016, 88 (22), 11084–11091. https://doi.org/10.1021/acs.analchem.6b03091.

(26) Zhou, Z.; Tu, J.; Xiong, X.; Shen, X.; Zhu, Z.-J. LipidCCS: Prediction of Collision Cross-Section Values for Lipids with High Precision To Support Ion Mobility–Mass Spectrometry-Based Lipidomics. Anal. Chem. 2017, 89 (17), 9559–9566. https://doi.org/10.1021/acs.analchem.7b02625.

(27) Plante, P.-L.; Francovic-Fontaine, É.; May, J. C.; McLean, J. A.; Baker, E. S.; Laviolette, F.; Marchand, M.; Corbeil, J. Predicting Ion Mobility Collision Cross-Sections Using a Deep Neural Network: DeepCCS. Anal Chem 2019, 91 (8), 5191–5199. https://doi.org/10.1021/acs.analchem.8b05821.

(28) Gabelica, V.; Shvartsburg, A. A.; Afonso, C.; Barran, P.; Benesch, J. L. P.; Bleiholder, C.; Bowers, M. T.; Bilbao, A.; Bush, M. F.; Campbell, J. L.; Campuzano, I. D. G.; Causon, T.; Clowers, B. H.; Creaser, C. S.; De Pauw, E.; Far, J.; Fernandez-Lima, F.; Fjeldsted, J. C.; Giles, K.; Groessl, M.; Hogan, C. J.; Hann, S.; Kim, H. I.; Kurulugama, R. T.; May, J. C.; McLean, J. A.; Pagel, K.; Richardson, K.; Ridgeway, M. E.; Rosu, F.; Sobott, F.; Thalassinos, K.; Valentine, S. J.; Wyttenbach, T. Recommendations for Reporting Ion Mobility Mass Spectrometry Measurements. Mass Spectrom Rev 2019, 38 (3), 291–320. https://doi.org/10.1002/mas.21585.

(29) Forsythe, J. G.; Petrov, A. S.; Walker, C. A.; Allen, S. J.; Pellissier, J. S.; Bush, M. F.; Hud, N. V.; Fernández, F. M. Collision Cross Section Calibrants for Negative Ion Mode Traveling Wave Ion Mobility-Mass Spectrometry. Analyst 2015, 140 (20), 6853–6861. https://doi.org/10.1039/C5AN00946D.

(30) Hines, K. M.; Ross, D. H.; Davidson, K. L.; Bush, M. F.; Xu, L. Large-Scale Structural Characterization of Drug and Drug-Like Compounds by High-Throughput Ion Mobility-Mass Spectrometry. Anal Chem 2017, 89 (17), 9023–9030. https://doi.org/10.1021/acs.analchem.7b01709.

(31) Li, A.; Conant, C. R.; Zheng, X.; Bloodsworth, K. J.; Orton, D. J.; Garimella, S. V. B.; Attah, I. K.; Nagy, G.; Smith, R. D.; Ibrahim, Y. M. Assessing Collision Cross Section Calibration Strategies for Traveling Wave-Based Ion Mobility Separations in Structures for Lossless Ion Manipulations. Anal Chem 2020, 92 (22), 14976–14982. https://doi.org/10.1021/acs.analchem.0c02829.

(32) Ridenour, W. B.; Kliman, M.; McLean, J. A.; Caprioli, R. M. Structural Characterization of Phospholipids and Peptides Directly from Tissue Sections by MALDI Traveling-Wave Ion Mobility-Mass Spectrometry. Anal Chem 2010, 82 (5), 1881–1889. https://doi.org/10.1021/ac9026115.

