## Supplemental Information for "Ion Mobility for Unknown Metabolite Identification: Hope or Hype?"

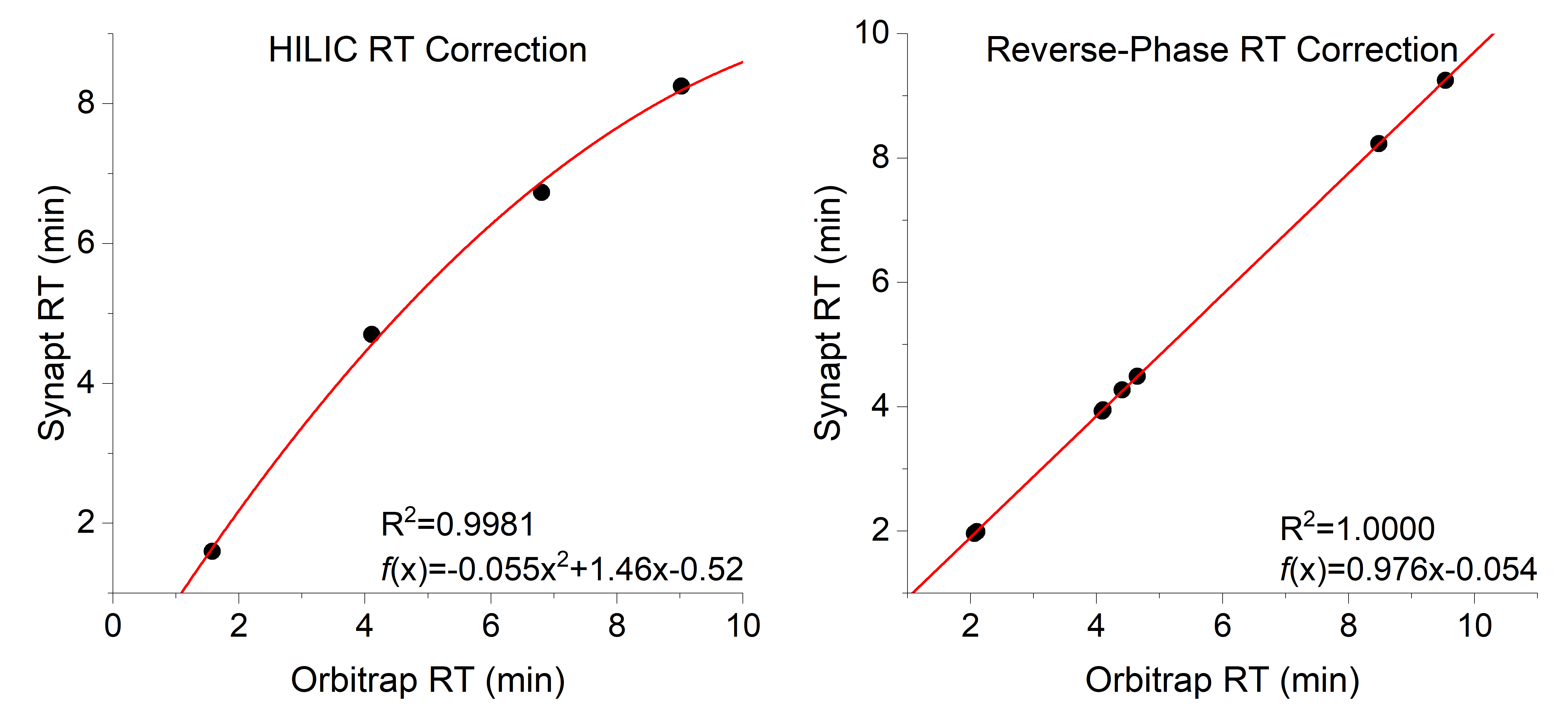

Figure S1. Minor chromatography differences between the LC-MS and LC-IM-MS platforms used necessitated retention time (RT) correction in order to correctly match features between runs. Shown are the correction curves for all standards as measured in the positive ion polarity, with the RT for both platforms plotted on each axis. A polynomial correction formula was used for the HILIC data and a linear correction formula for reverse-phase data.

**Ranked ANOVA-Additional Details**

To account for variation in the amplitude in the chromatograms across batches a normalization process is typically applied. The most common is a total ion chromatograph correction. This approach has several well-known issues. Our goal in this first feature selection step is simply to identify features with large and consistent differences between the mutant and control. To this end we elected to use rank transformation with 1,000 bins followed by an ANOVA as illustrated in Figure S2. The ranking approach assigns high values to relatively intense features within a chromatogram and low values to features with low intensity. By fixing the number of bins, small differences in the number of features identified in the chromatogram are minimized. The ANOVA is as a screening procedure that will identify features that are consistently high in one group and consistently low in the other group. Here we are interested in identifying a subset of features for further elucidation, and so we elected to use a nominal p-value of 0.05 to minimize the type II error, as is typical of sequential screening studies. Residuals were examined and behaved according to model assumptions. We compared ranked unbinned to rank binned with several different numbers of bins. Results were nearly identical between the 1,000 bins and the rank transformation alone. The results from 1,000 bins were slightly more conservative than the unbinned ranked data in the number of significant features.

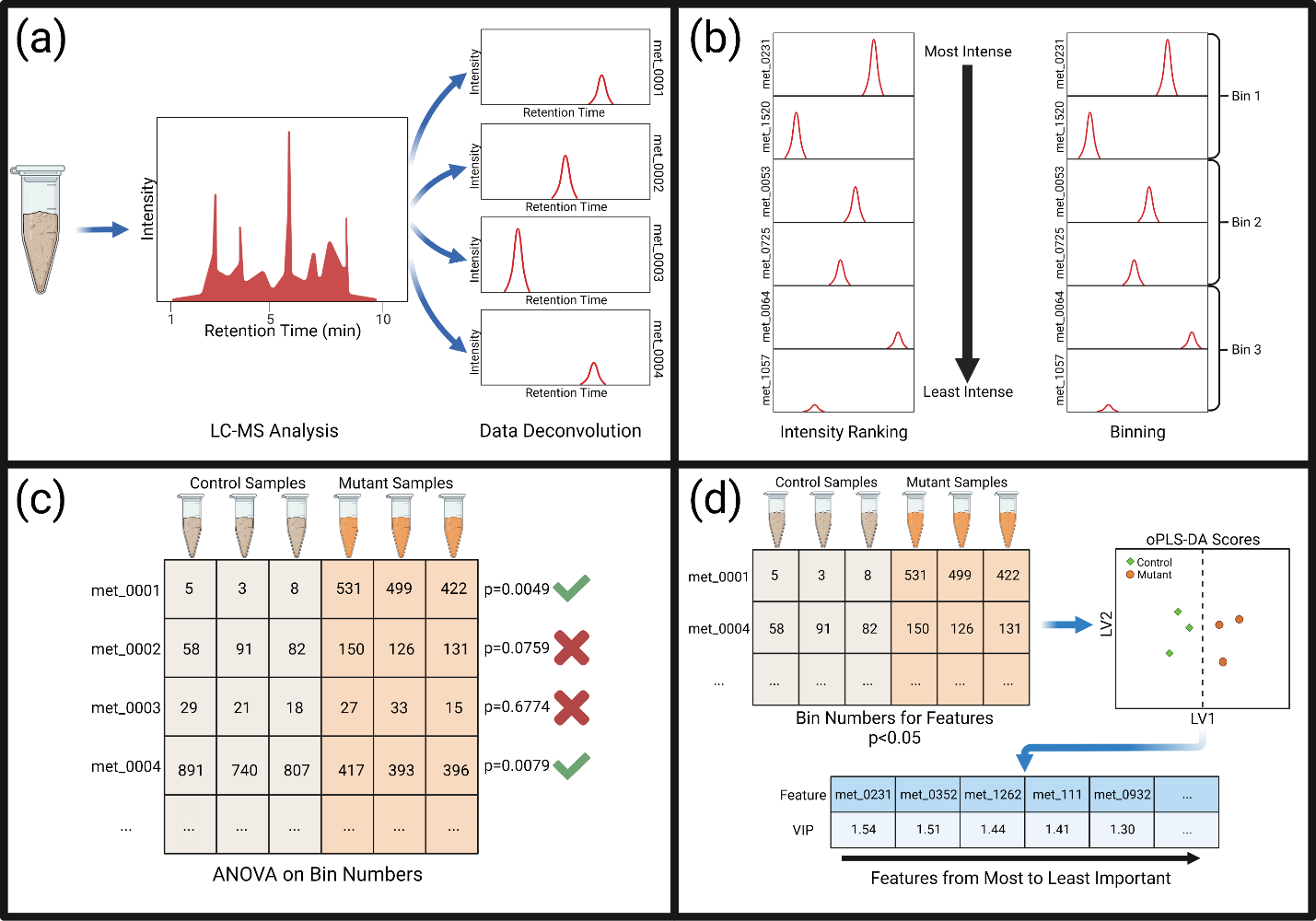

**Figure S2**: Data processing of LC-MS data acquired via Thermo Orbitrap ID-X began with data deconvolution within Thermo Compound Discoverer 3.1 to produce a list of ion features and their corresponding peak areas (a). These features were ranked from most intense to least intense within each sample, and then binned into 1000 bins, giving each feature a bin number for each sample representative of its relativity intensity in that sample (b). ANOVA was performed between the mutant and control strain sample bins to identify features that were significantly different (p<0.05) (c). Significant features based on ANOVA were retained, autoscaled and used to build oPLS-DA models, yielding VIP scores for each feature (d). Features with the top VIP scores were subject to structural annotation.

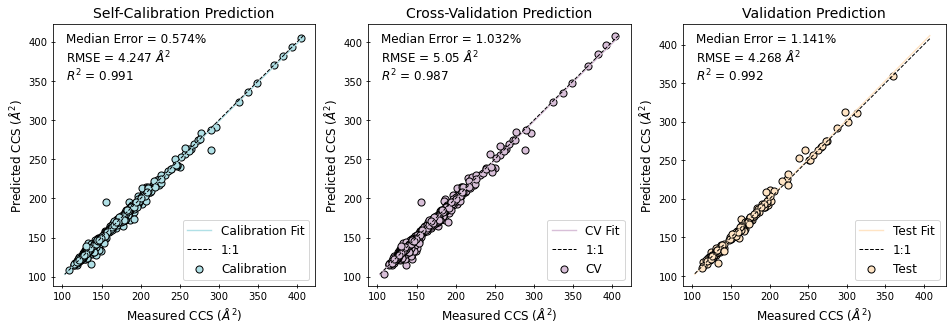

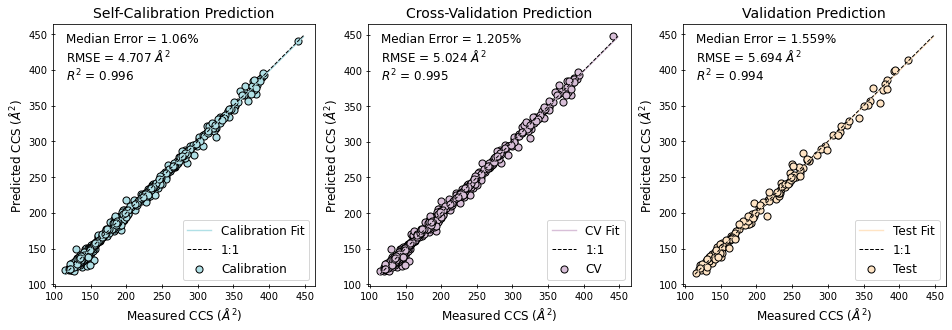

Figure S3. Validation results for CCSP 2.0 for [M-H]^-^ ions (top) and [M+H]^+^ ions (bottom). Self-calibration shows the errors of the predicted CCS for all entries in the training set. Cross-validation errors results from leaving out 20% of the training data and constructing the model from the remaining 80%. External validation was performed with a separate test set as shown in the validation prediction panel.

**Figure S4**. Variable Importance in Projection (VIP) scores vs. LV1 oPLS-DA score for feature HN_196 is shown as a green point in (a) demonstrating the high importance for this feature in discriminating mutant strain RB2347 from control strain PD1074. The hierarchy in (b) shows the reduction of all compounds with a matching elemental formula, to those generated from in silico MS/MS prediction, to those MS/MS candidates within the ±3% CCS error range. Top ranked candidate structures from SIRIUS 4 are shown in (c) with their predicted vs. measured CCS error. The MS/MS spectrum for this feature is compared against the MS/MS spectrum acquired from a pure standard of N-Acetylaspartate, the rank 1 MS/MS candidate structure (d). This is repeated for features HN_271 and HN_480 in mutant strain VC1265 in panels (e-h) and (i-l), respectively.

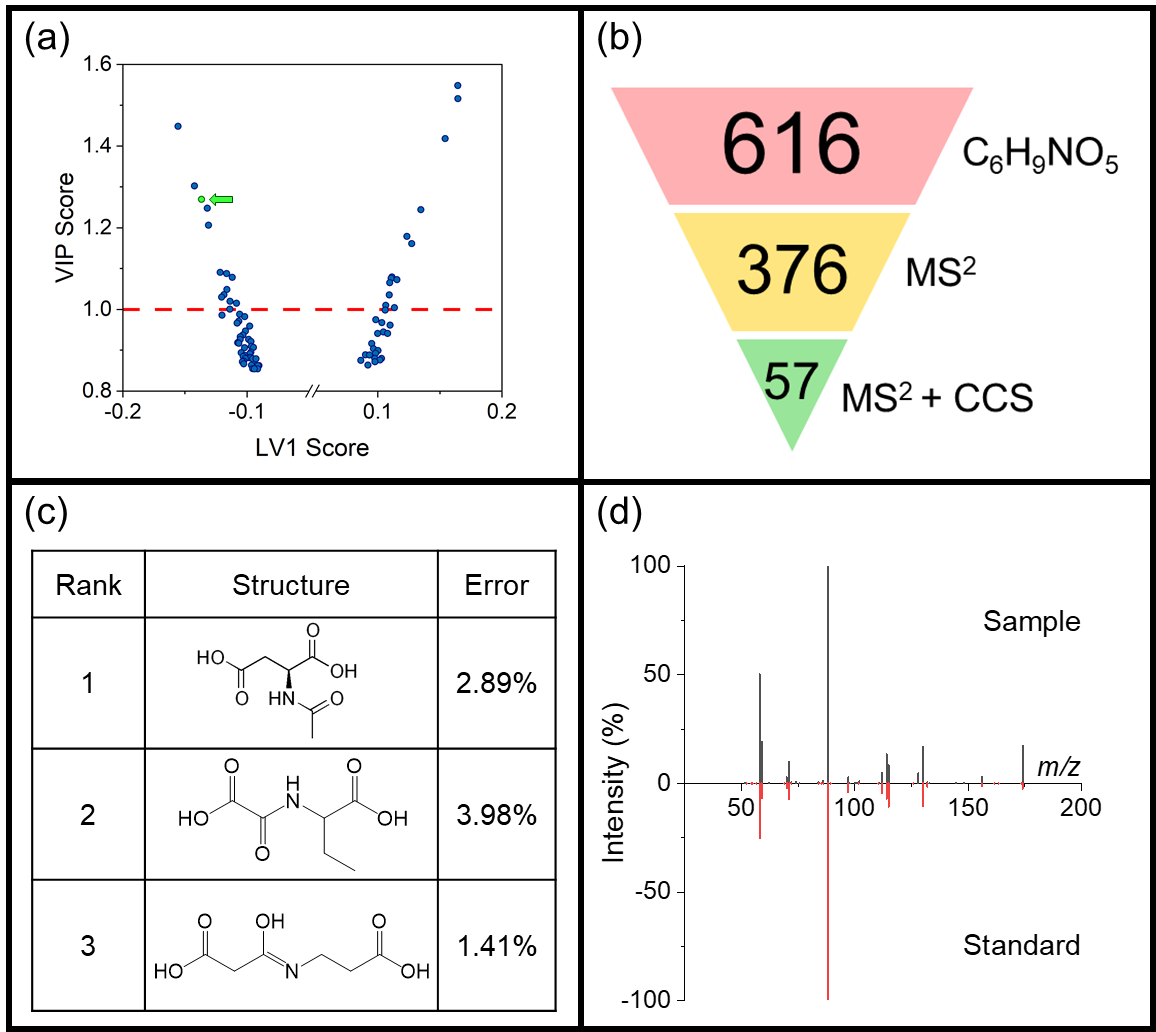

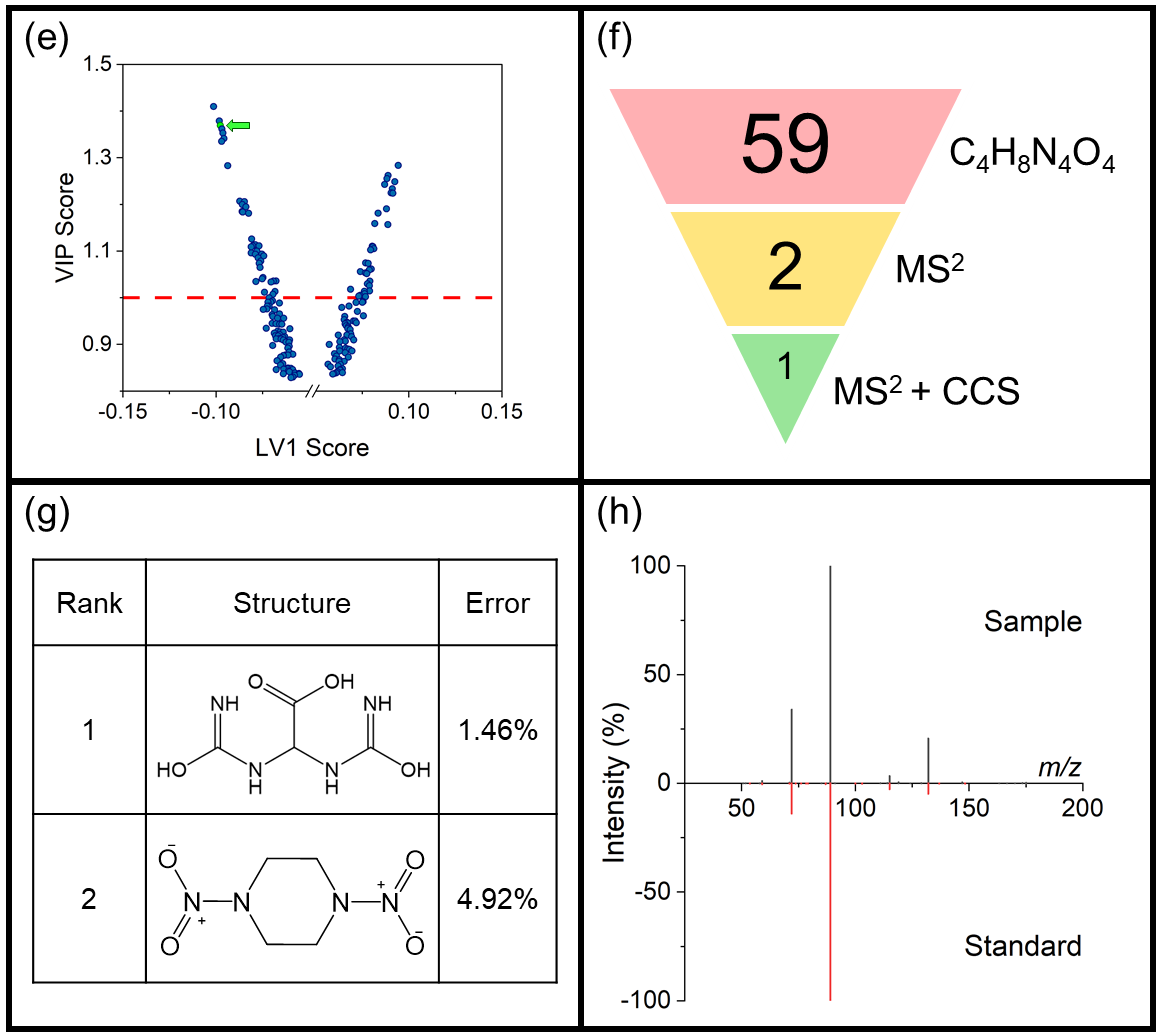

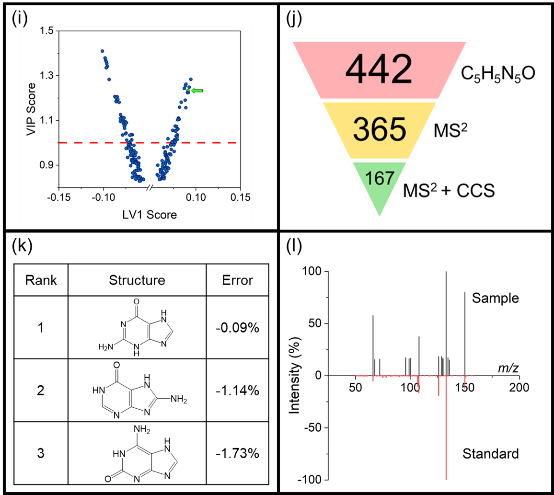

**Table S1**. Internal standards used for retention time alignment in polar and non-polar extracts.

| **Standard Name** | **Formula** | **Fraction** |
| --- | --- | --- |
| L-Arginine (^13^C_6_, 99%) | [13]C_6_H_14_N_4_O_2_ | Polar |
| Hippuric Acid (Benzoyl-d_5_, 98%) | C_9_H_4_D_5_NO_3_ | Polar |
| Hypoxanthine (^13^C_5_, 99%) | [13]C_5_H_4_N_4_O | Polar |
| L-Methionine (1-^13^C, 99%; methyl-d_3_, 98%) | [13]CC_4_H_8_D_3_NO_2_S | Polar |
| 15:0-18:1(d_7_) PC | C_41_H_73_D_7_NO_8_P | Non-polar |
| 18:1(d_7_) Lyso PC | C_26_H_45_D_7_NO_7_P | Non-polar |
| 15:0-18:1(d_7_) PE | C_38_H_67_D_7_NO_8_P | Non-polar |
| 18:1(d_7_) Lyso PE | C_23_H_39_D_7_NO_7_P | Non-polar |
| 15:0-18:1(d_7_) PG | C_39_H_68_D_7_O_10_P_2_ | Non-polar |
| 15:0-18:1(d_7_) PI | C_42_H_72_D_7_O_13_P | Non-polar |
| 15:0-18:1(d_7_) PS | C_39_H_67_D_7_NO_10_P | Non-polar |
| 15:0-18:1(d_7_)-15:0 TAG | C_51_H_89_D_7_O_6_ | Non-polar |
| 15:0-18:1(d_7_) DAG | C_36_H_61_D_7_O | Non-polar |
| 18:1(d_7_) Chol Ester | C_45_H_71_D_7_O_2_ | Non-polar |
| d18:1-18:1(d_9_) SM | C_41_H_72_D_9_N_2_O_6_P | Non-polar |
| Cholesterol-d_7_ | C_27_H_39_D_7_O | Non-polar |

**Table S2**. Chromatography gradient for HILIC chromatography. Solvent A was 80:20 water:acetonitrile with 0.1% formic acid and 10 mM ammonium formate. Solvent B was 0.1% formic acid in H_2_O.

| **Time** | **Flow (mL/min)** | **%B** |
| --- | --- | --- |
| 0 | 0.4 | 95 |
| 0.5 | 0.4 | 95 |
| 8 | 0.4 | 40 |
| 9.4 | 0.4 | 40 |
| 9.5 | 0.4 | 95 |
| 11 | 0.4 | 95 |
| 12 | 0.4 | 95 |

**Table S3**. Chromatography gradient for HILIC chromatography. Solvent A was 40:60 water:acetonitrile with 0.1% formic acid and 10 mM ammonium formate. Solvent B was 90:10 isopropanol:acetonitrile with 0.1% formic acid and 10 mM ammonium formate.

| **Time** | **Flow (mL/min)** | **%B** |
| --- | --- | --- |
| 0 | 0.4 | 20 |
| 1 | 0.4 | 60 |
| 5 | 0.4 | 70 |
| 5.5 | 0.4 | 85 |
| 8 | 0.4 | 90 |
| 8.2 | 0.4 | 100 |
| 10.5 | 0.4 | 100 |
| 10.7 | 0.4 | 20 |
| 12 | 0.4 | 20 |

**Table S4**. Full Synapt G2-S instrument parameters. Parameters not listed were left at their automatic values.

| **Tab** | **Name** | **Setting** |
| --- | --- | --- |
| ES+ | Capillary | 3.00 |
| ES+ | Sampling cone | 30 |
| ES+ | Source offset | 40 |
| ES+ | Source temp | 80 |
| ES+ | Desolvation temp | 250 |
| ES+ | Cone gas flow | 50 |
| ES+ | Desolvation gas flow | 650 |
| ES+ | Nebulizer gas flow | 7.0 |
| Stepwave | SW1 wave height | 5 |
| Stepwave | SW1 wave velocity | 300 |
| Stepwave | SW2 wave height | 5 |
| Stepwave | SW2 wave velocity | 300 |
| Stepwave | SW2 offset | 15 |
| Stepwave | Stepwave RF | 150 |
| Stepwave | Ion guide RF | 200 |
| Instrument | IMS gas flow | 70 |
| Triwave | IMS wave velocity | 600-200 (ramp full cycle) |
| Triwave | IMS wave height | 22 |
| Triwave DC | Trap entrance | 3 |
| Triwave DC | Trap bias | 30 |
| Triwave DC | Trap DC | -4 |
| Triwave DC | Trap exit | 0 |
| Triwave DC | IMS entrance | 12 |
| Triwave DC | IMS He cell DC | 35 |
| Triwave DC | IMS He exit | -12 |
| Triwave DC | IMS bias | 10 |
| Triwave DC | IMS exit | 0 |

**Table S5**. oPLS-DA parameters within PLS toolbox 8.9.1 (Eigenvector Research, Inc.).

| This is a model of type | oPLSDA |
| --- | --- |
| Developed | 24-Jun-2021 |
| X-block | RB2347_HILICneg_sig05_with validation.xlsx 14 by 17 |
| Included | [ 1-14 ] [ 4 6 17-18 20-21 27-28 31-32 49 57-58 60 62 70 78 ] |
| Preprocessing | Autoscale |
| Y-block | y 14 by 2 |
| Included | [ 1-14 ] [ 1-2 ] |
| Preprocessing | Autoscale |
| Num. LVs | 2 |

**Table S6**. Parameters for genetic algorithm used for feature selection within oPLS-DA model.

| **Parameter** | **Setting** |
| --- | --- |
| Population size | 64 |
| Window width | 1 |
| % Initial terms | 30 |
| Target as % of variables | 0-30 |
| Penalty slope | 0.01 |
| Max generations | 200 |
| % at convergence | 50 |
| Mutation rate | 0.005 |
| Crossover | Double |
| Regression choice | PLS |
| #LVs | 14 |
| Cross-validation parameters | Contiguous |
| # of splits | 7 |
| # of iterations | 5 |
| Replicate runs | 2 |

**Table S7**. Description of failure points for features with both CCS and MS/MS data which could not complete the candidate structure filtering workflow.

| **Fraction** | **Polarity** | **RT (min)** | **Molecular Weight** | **Error** |
| --- | --- | --- | --- | --- |
| Non-polar | +ive | 4.379 | 745.644 | Insufficient training data for adduct |
| Non-polar | +ive | 5.189 | 773.675 | Insufficient training data for adduct |
| Non-polar | -ive | 5.702 | 1518.204 | CCS rollover |
| Non-polar | -ive | 3.341 | 1447.011 | Insufficient training data for adduct |
| Polar | +ive | 8.004 | 196.046 | No candidate structures from MS/MS |
| Polar | +ive | 9.025 | 395.098 | Insufficient training data for adduct |
| Polar | +ive | 7.822 | 514.135 | Insufficient training data for adduct |
| Polar | +ive | 4.190 | 425.226 | Insufficient training data for adduct |
| Non-Polar | -ive | 6.296 | 641.596 | No candidate structures from MS/MS |
| Non-Polar | -ive | 2.971 | 649.469 | Poor arrival time assignment |
| Non-Polar | -ive | 2.965 | 699.484 | Poor arrival time assignment |
| Non-Polar | -ive | 6.781 | 669.628 | Poor arrival time assignment |
| Polar | +ive | 8.298 | 500.119 | Poor arrival time assignment |
| Polar | -ive | 6.560 | 342.071 | Insufficient training data for adduct |
| Polar | -ive | 2.625 | 215.094 | No candidate structures from MS/MS |
